## Supplemental Figures for "Single Cell RNA Sequencing of the Adult *Drosophila* Eye Reveals Distinct Clusters and Novel Marker Genes for All Major Cell Types"

Supp Figure 1. Monocle 3 clustering of 1-day, 3-day, and 7-day old adult eyes do not show any clear progression of transcriptome changes.

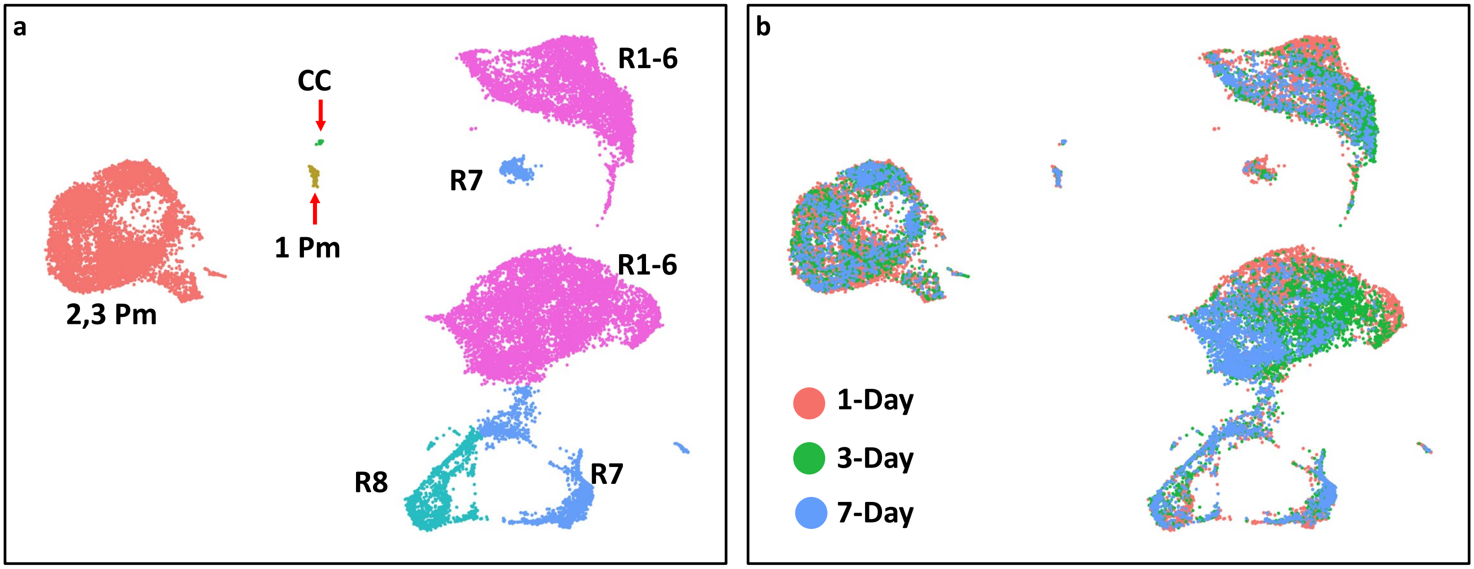

**Supplemental Figure 1. Monocle 3 clustering of 1-day, 3-day, and 7-day old adult eyes do not show any clear progression of transcriptome changes. a**) Monocle 3 clustering of 1-day, 3-day and 7-day old adult eyes showing the major cell types of the eye. **b**) Same cluster as in A but showing the sample age of each cell. All clusters show an intermixing of cells from the three time points.

Supp Figure 2. R8 FeaturePlots for 1-day and 3-day old adult eyes

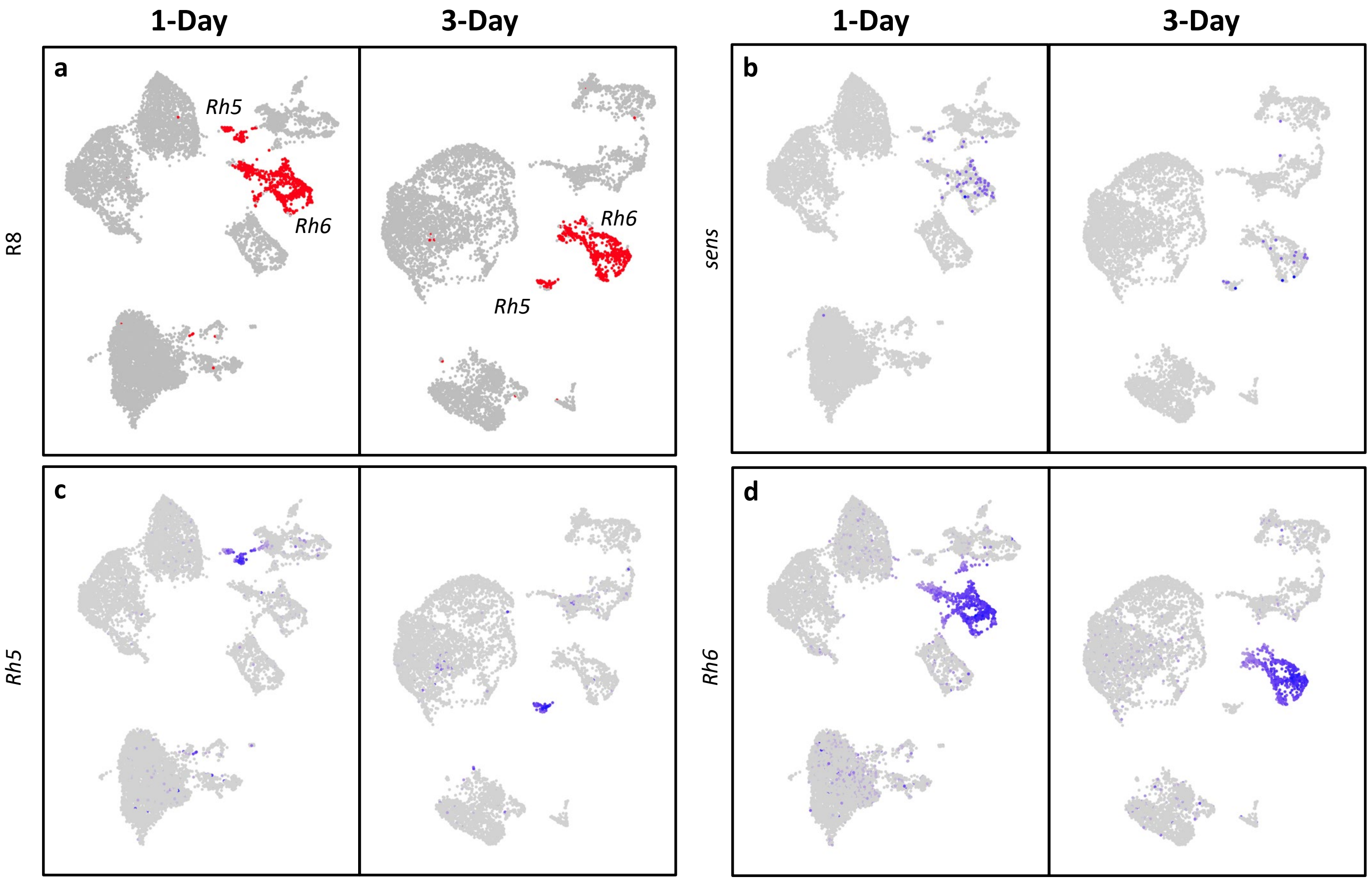


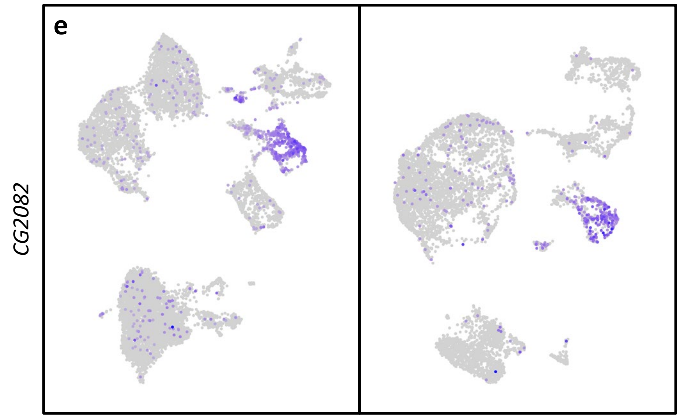


**f** **Numbers of *Rh5* and *Rh6* R8 cells**

| *Rh* | 1-Day | 3-Day | 7-Day | Expected |
|---|---|---|---|---|
| *Rh5* | 113 (14%) | 58 (10%) | 63 (13%) | ~30% |
| *Rh6* | 678 (86%) | 517 (90%) | 441 (88%) | ~70% |
| **Total** | **791** | **575** | **504** | **100%** |

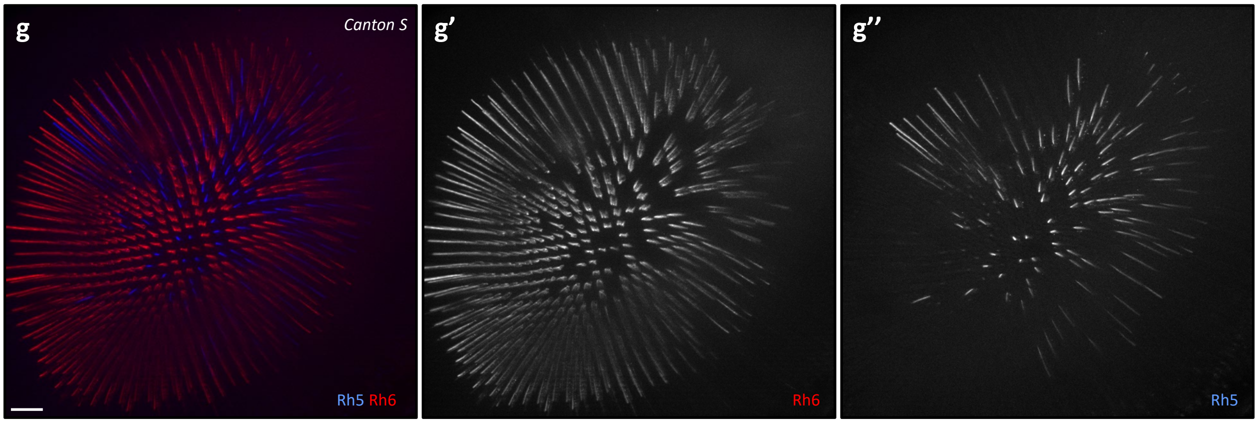

**Supplemental Figure 2. R8 FeaturePlots for 1-day and 3-day old adult eyes. a**) Cluster plots showing R8 cells in 1-day and 3-day eye data sets (red).**b-e**) FeaturePlots showing the expression of *sens* (**b**), *Rh5* (**c**), *Rh6* (**d**) and *CG2082* (**e**) in 1-day and 3-day old adult eyes. Cells expressing *sens* and *CG2082* are brought to the front. **f**) Table showing the numbers of *Rh5* and *Rh6* positive R8 cells. **g**) Immunostaining of a *CantonS* eye showing *Rh5* and *Rh6* expression at the expected ~30:70 ratio. Scale bar: 20 μm.

Supp Figure 3. R7 FeaturePlots for 1-day and 3-day old adult eyes

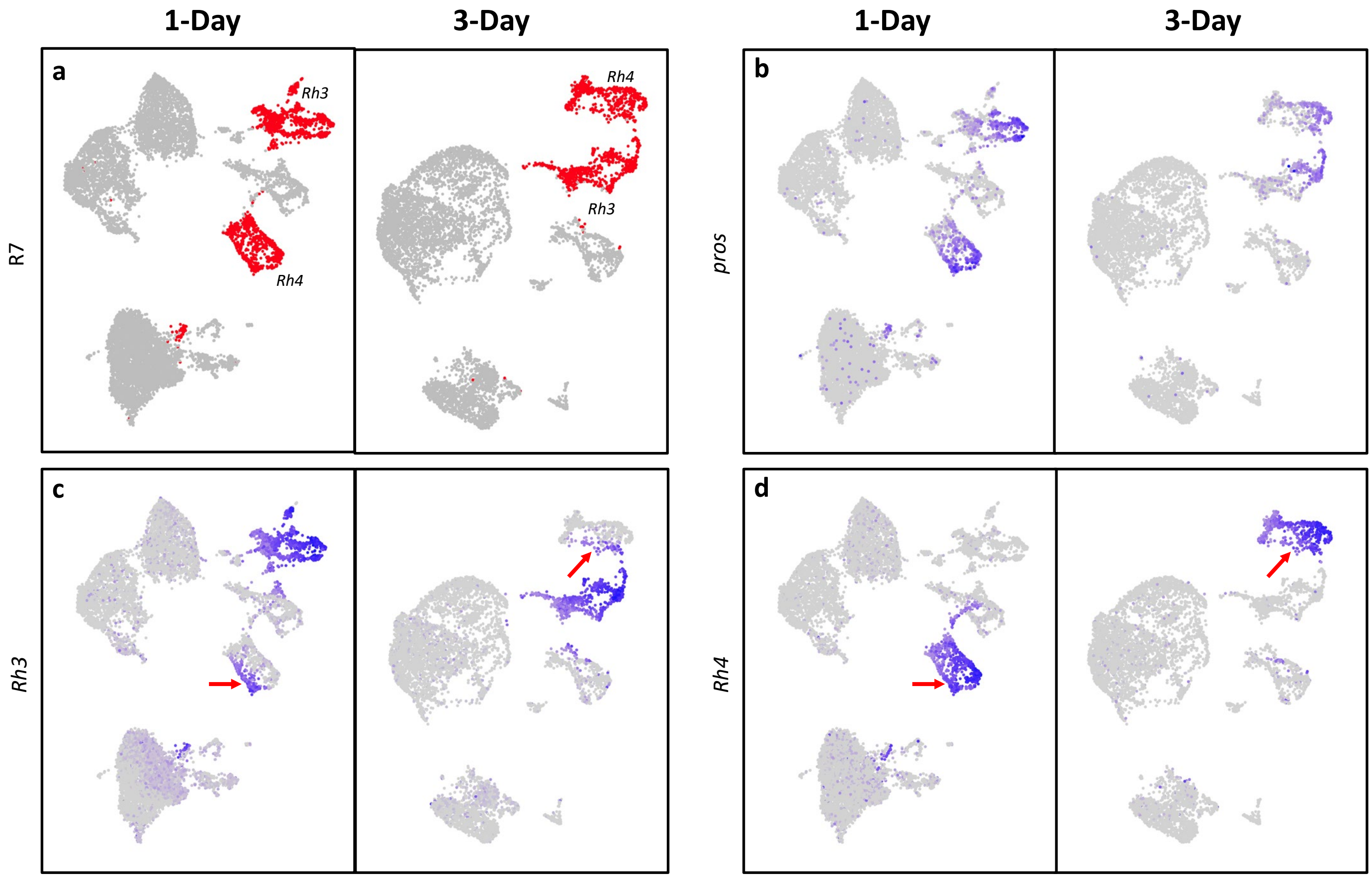


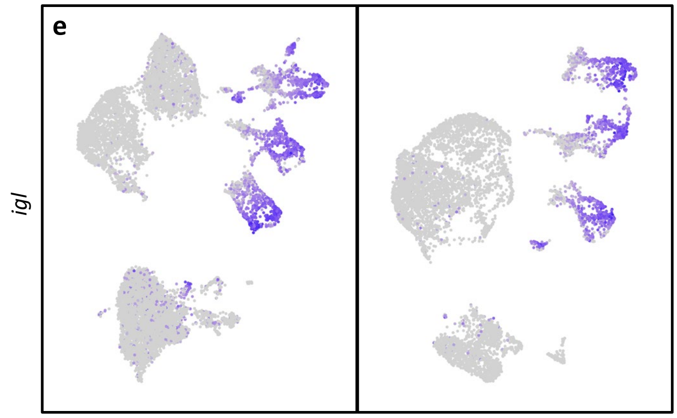


**f** **Numbers of Pale (*Rh3*) and Yellow (*Rh4*) R7 cells**

| Pale / Yellow | 1-Day | 3-Day | 7-Day | Expected |
|---|---|---|---|---|
| Pale | 967 (55%) | 666 (53%) | 525 (55%) | ~30% |
| Yellow | 803 (45%) | 585 (47%) | 428 (45%) | ~70% |
| **Total** | **1,770** | **1,251** | **953** | **100%** |

**Supplemental Figure 3. R7 FeaturePlots for 1-day and 3-day old adult eyes. a**) Cluster plots showing R7 cells in 1-day and 3-day eye data sets (red).**b-e**) FeaturePlots showing the expression of *pros* (**b**), *Rh3* (**c**), *Rh4* (**d**) and *igl* (**e**) in 1-day and 3-day old adult eyes. Cells expressing *pros* and *igl* are brought to the front. **f**) Table showing the numbers of *pale* and *yellow* positive R7 cells.

Supp Figure 4. Dorsal rim area FeaturePlots for 1-day, 3-day and 7-day old adult eyes

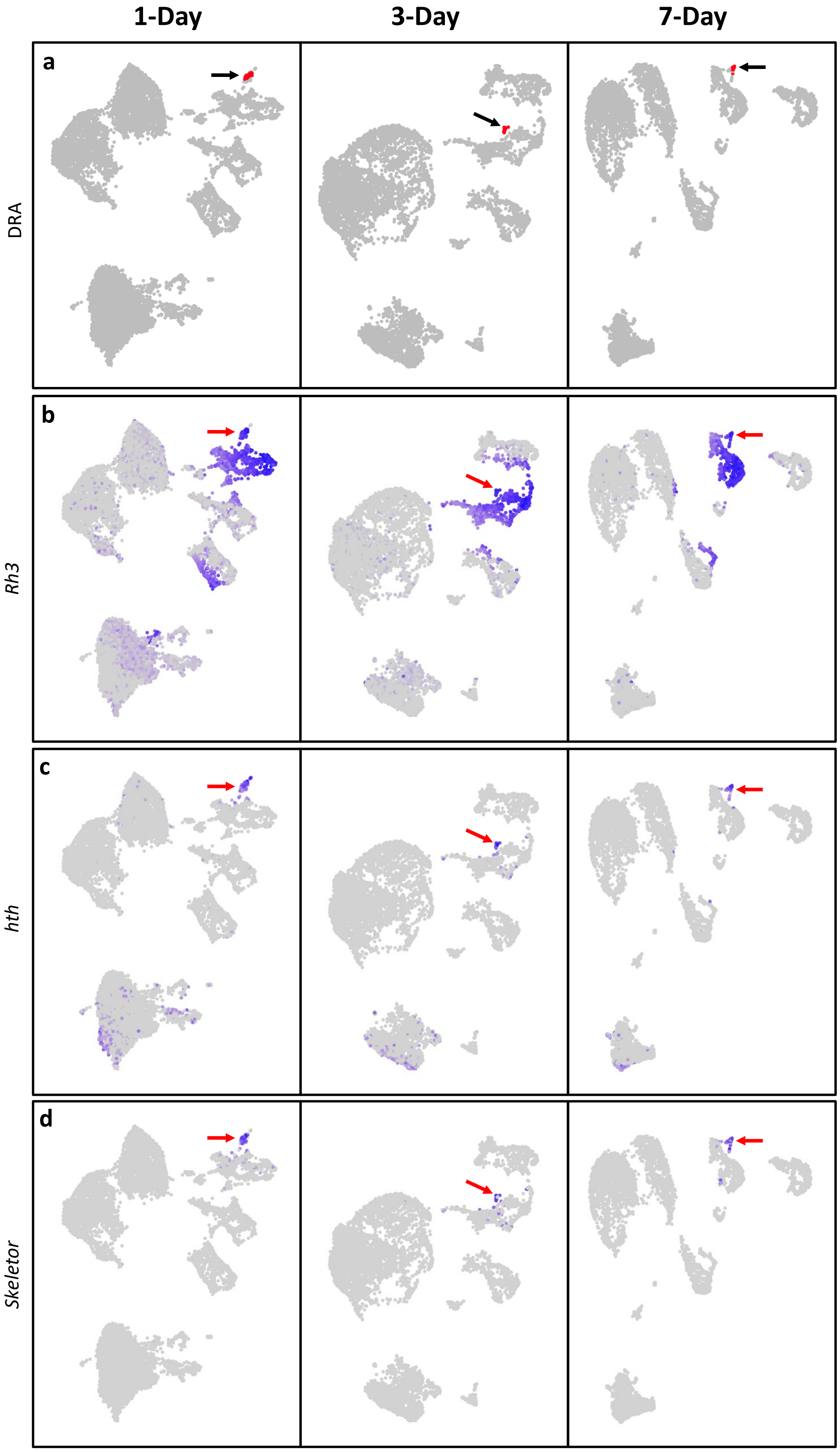

**Supplemental Figure 4. Dorsal rim area FeaturePlots for 1-day, 3-day and 7-day old adult eyes. a**) Cluster plots showing dorsal rim area R7/R8 cells in 1-day, 3-day and 7-day old eye data sets (red). Black arrow points to the dorsal rim area R7/R8. **b-d**) FeaturePlots showing the expression of *Rh3* (**b**), *hth* (**c**) and *Skeletor* (**d**) in 1-day, 3-day and 7-day old adult eyes.

Supp Figure 5. R1-6 cluster and feature plots for 1 day and 3 days old adult eyes

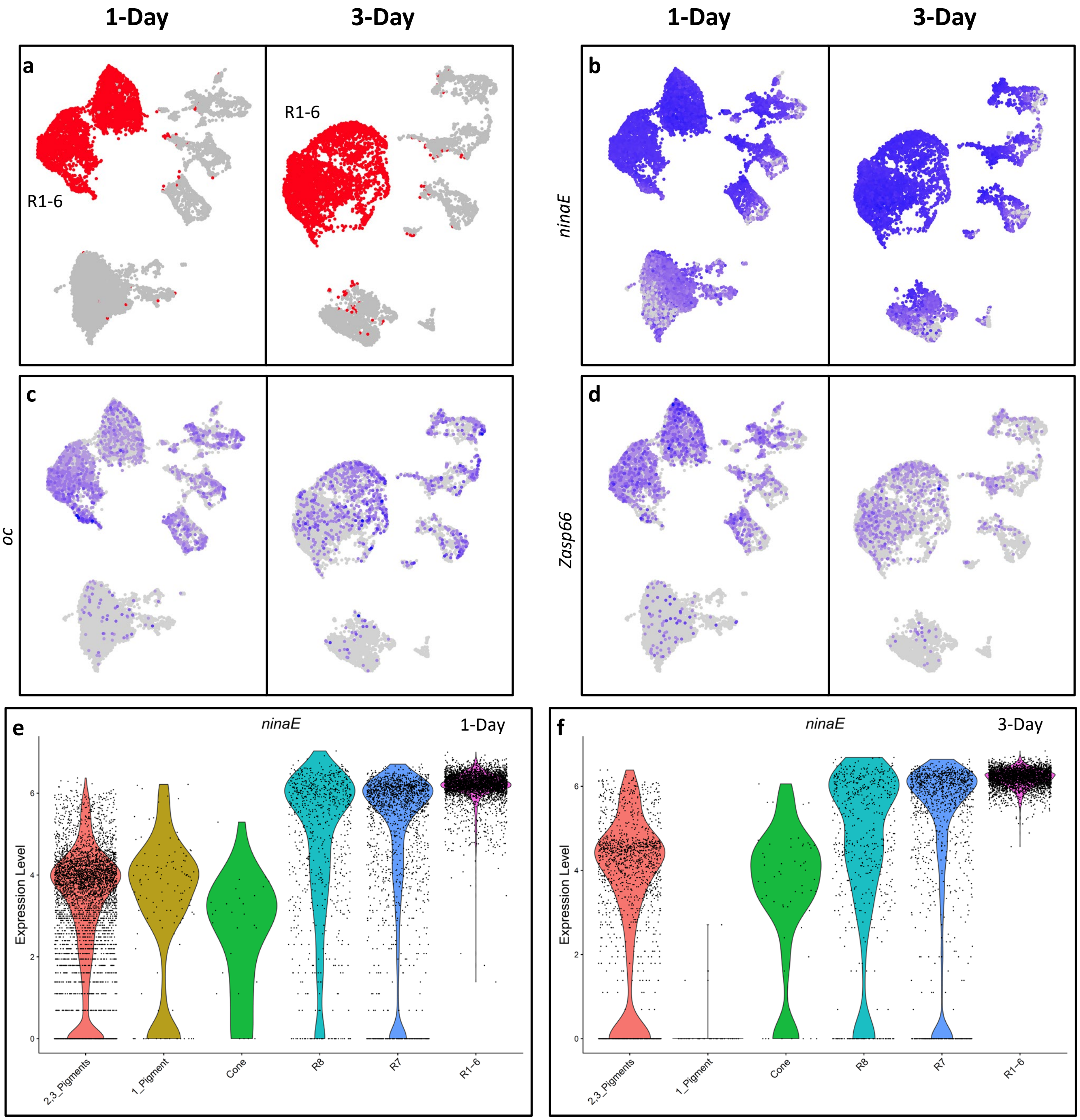

**Supplemental Figure 5. R1-6 FeaturePlots for 1-day and 3-day old adult eyes. a**) Cluster plots showing R1-6 cells in 1-day and 3-day eye data sets (red). **b-d**) FeaturePlots showing the expression of *ninaE* (**b**), *oc* (**c**), and *Zasp66* (**d**) in 1-day and 3-day old adult eyes. Cells expressing *oc* and *Zasp66* are brought to the front. **e**, **f**) Violin plots showing *ninaE* expression in 1-day (**e**) and 3-day (**f**) old adult eyes

Supp Figure 6. Cone cell FeaturePlots for 1-day and 3-day old adult eyes

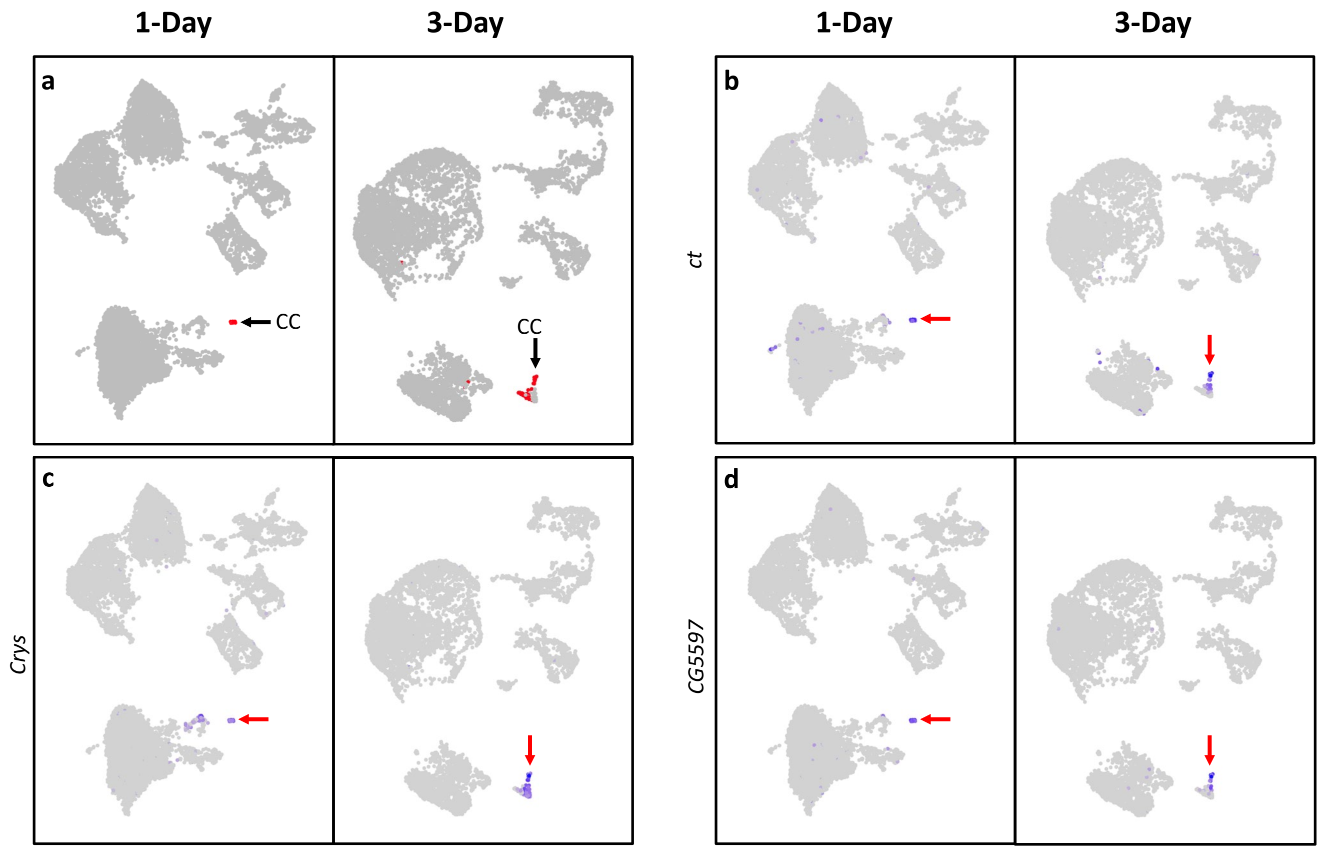

**Supplemental Figure 6. Cone cell FeaturePlots for 1-day and 3-day old adult eyes. a**) Cluster plots showing cone cells in 1-day and 3-day eye data sets (red). Black arrows point to the cone cell clusters. **b-d**) FeaturePlots showing the expression of *ct* (**b**), *Crys* (**c**), *CG5597* (**d**) in 1-day and 3-day old adult eyes. Cells expressing *ct*, *Crys* and *CG5597* are brought to the front.

Supp Figure 7. Pigment cell FeaturePlots for 1-day and 3-day old adult eyes

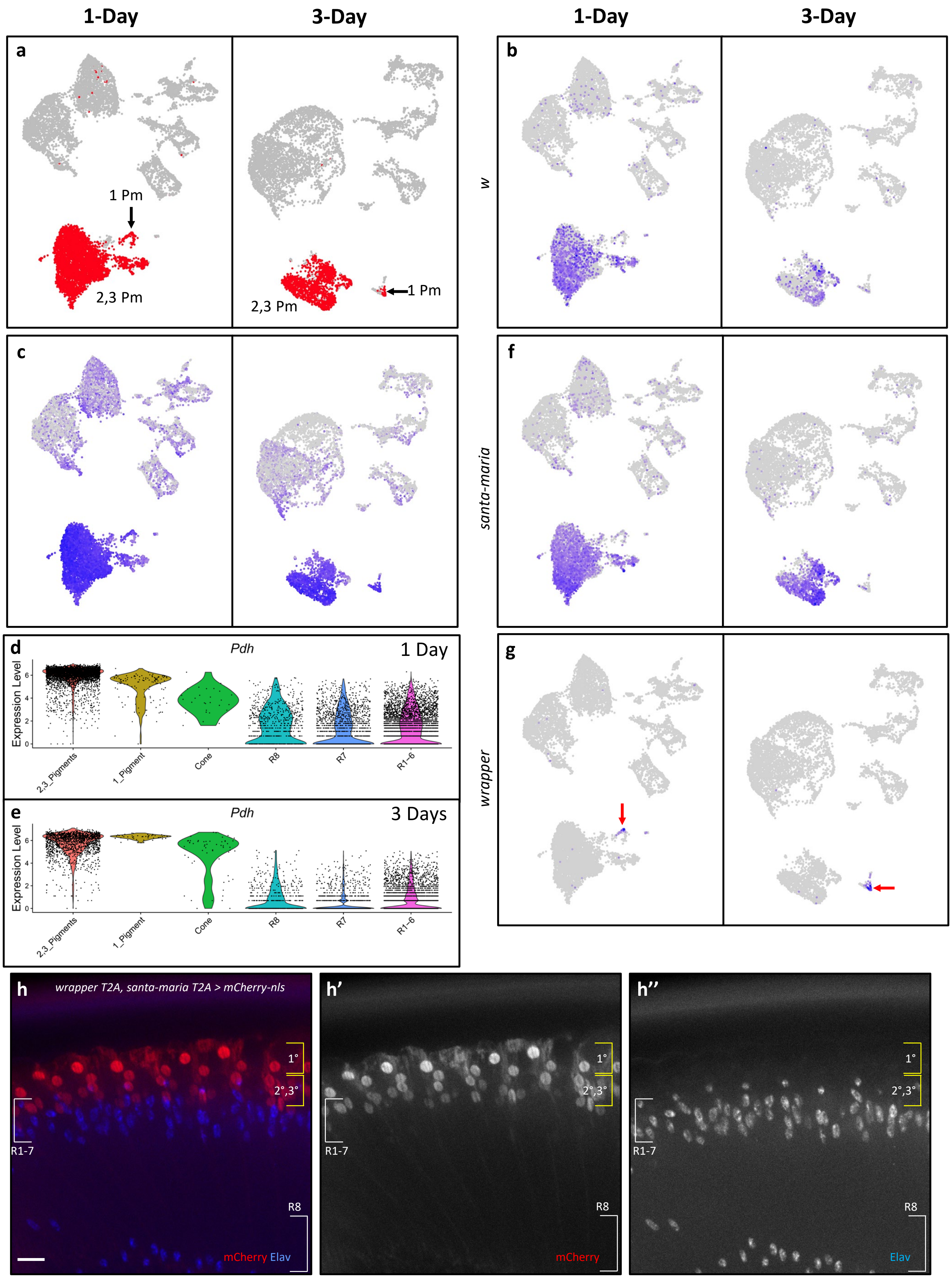

**Supplemental Figure 7. Pigment cell FeaturePlots for 1-day and 3-day old adult Eyes. a**) Cluster plots showing pigment cells cells in 1-day and 3-day eye data sets (red). Black arrows point to the smaller primary pigment cell cluster (1 Pm); the remaining red cells are secondary and tertiary pigment cells (2,3 Pm). **b**, **c**) FeaturePlots showing the expression of *w* (**b**) and *Pdh* (**c**) in 1-day and 3-day old adult eyes. **d**, **e**) Violin plots showing *Pdh* expression in 1-day (**d**) and 3-day (**e**) old adult eyes. **f**, **g**) FeaturePlots showing the expression of *santa-maria* (**f**) and *wrapper* (**g**) in 1-day and 3-day old adult eyes. Red arrows point to *wrapper* positive cells in **g**. **h**) Coronal view of a *wrapper-T2A-Gal4, santa-maria-T2A-Gal4* > *UAS-mCherry-nls* adult eye with mCherry stained red and Elav stained blue. Scale bar: 10 μm.

Supp Figure 8. R7 clustering is not affected by the removal of *Rh5/6* in adult eye clustering

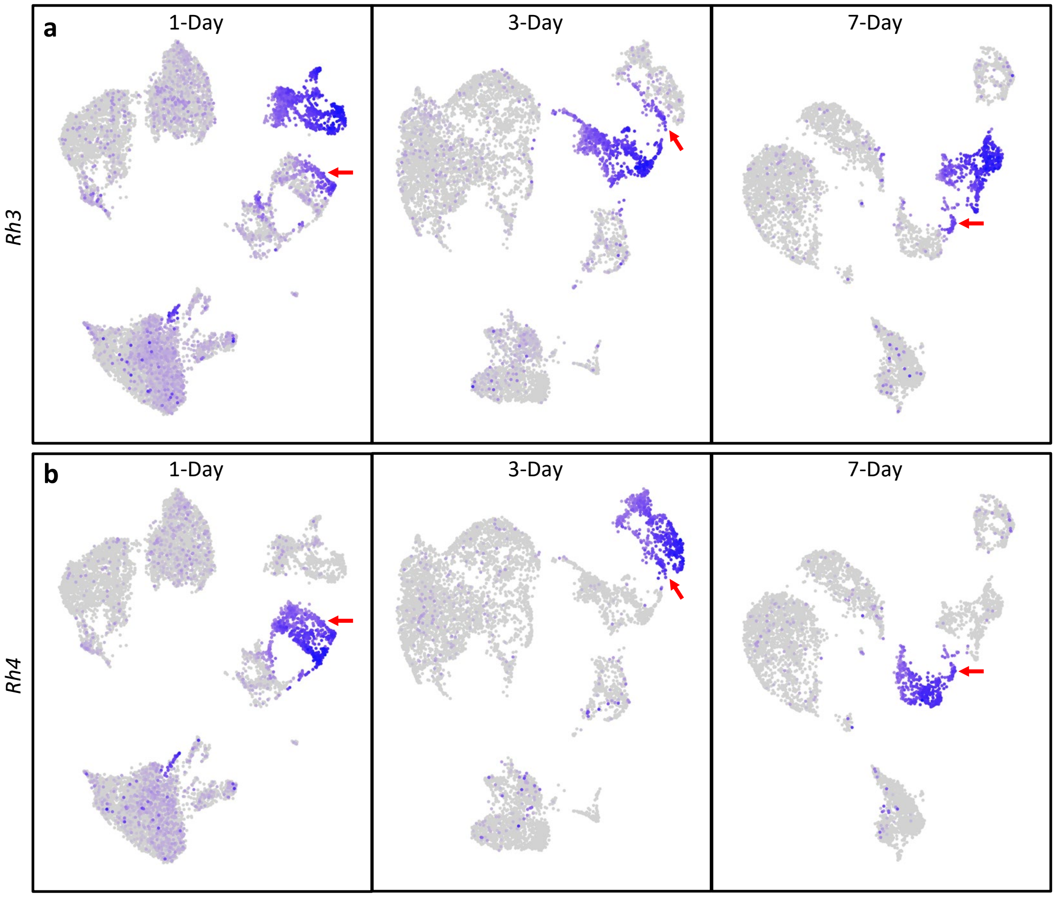

**Supplemental Figure 8. R7 clustering is not affected by the removal of *Rh5/6* in adult eye clustering.**

**a**) FeaturePlots of *Rh3* of 1-day, 3-day and 7-day old adult eye data where *Rh5*/6 count data were removed prior to clustering. **b**) FeaturePlots of *Rh4* of 1-day, 3-day and 7-day old adult eye data where *Rh5*/6 count data were removed prior to clustering. Red arrows point to the dorsal third R7s where *Rh3* and *Rh4* are co-expressed. Cells expressing *Rh3* and/or *Rh4* are brought to the front.
